# Bioenergetic signature of Synaptic mitochondria

**DOI:** 10.1101/2024.11.28.625855

**Authors:** Andreia Faria-Pereira, Vanessa A. Morais

**Author notes:** CORRESPONDING AUTHOR Vanessa A. Morais.

## Abstract

Synaptic transmission is the most energy-demanding processes in the brain and here we show that mitochondria have developed specific properties to efficiently support neurotransmission. It is a known fact that mitochondria at synapses need to be able to deal with a dynamic range of energetic needs overtime and to adapt between resting and high stimulation conditions. However, how mitochondria are adjusting to this requirement was not yet clear. Here, we show that synaptic mitochondria have a distinct bioenergetic profile presenting a stronger ability to respond to respiratory stimulus. These features are explained by a dichotomic Complex I activity pattern where synaptic mitochondria present a decreased enzymatic activity of individual Complex I, yet mitochondria at synapse present a dramatically enhanced Complex I+III combined activity. These bioenergetics features may endow synaptic mitochondria with the necessary mechanisms to adapt to the flexible bioenergetic environment present at synapses.

## INTRODUCTION

Neurons are high energy demanding cells with a unique morphological polarization. As ATP has a low diffusion rate^1^, neuronal energetic demands need to be met *in loco* for each neuronal sub-compartment. Synaptic transmission is one of the most energy-demanding processes in neurons^2–5^, and oxygen consumption promptly increases in response to stimulation^6,7^. These synaptic energetic requirements are mainly fulfilled by mitochondrial oxidative phosphorylation which under sustained synaptic activity is complemented by glycolysis^4,8–12^. Due to the unique intracellular environment, one could postulate that mitochondria at synapses may have an “adapted” profile in comparison to mitochondria that reside in other parts of the neuron or brain. In line with this, lipidomic^13^ and proteomics^14,15^ studies show differences in the composition of synaptic mitochondria. Additionally, synaptic mitochondria have a distinct volume, size and ultrastructure compared to mitochondria elsewhere in the neuron and brain (non-synaptic mitochondria)^16–19^. Recent studies show that synaptic stimulation impacts mitochondrial dynamics and biogenesis, either through the recruitment of axonal mitochondria to synapses^20^, or through the approximately one-fourth increased synthesis of mitochondrial related proteins at synapses^21^. Additionally, these protein translation processes at post-synaptic sites appear to be fuelled by mitochondria^22^, emphasizing the interdependence between mitochondria function and sustained synaptic activity.

In 1989, Lai and Clark proposed that mitochondria in the brain have some degree of “microheterogeneity”^23^, however, were unable to clarify this heterogeneity at the molecular level. Here, we unveil bioenergetic adaptations of mitochondria at the synapse to meet their energetic requirements. Our results show that under resting conditions mitochondria have tailor-made “energetic machinery” on-hold to help responding to the higher energetic demands upon synaptic stimulation.

## RESULTS and DISCUSSION

### Isolation and morphological assessment of synaptic mitochondria

Synaptic and non-synaptic mitochondria were isolated from 8-week old mice through differential centrifugation steps (Figure 1a).

**Fig. 1.**
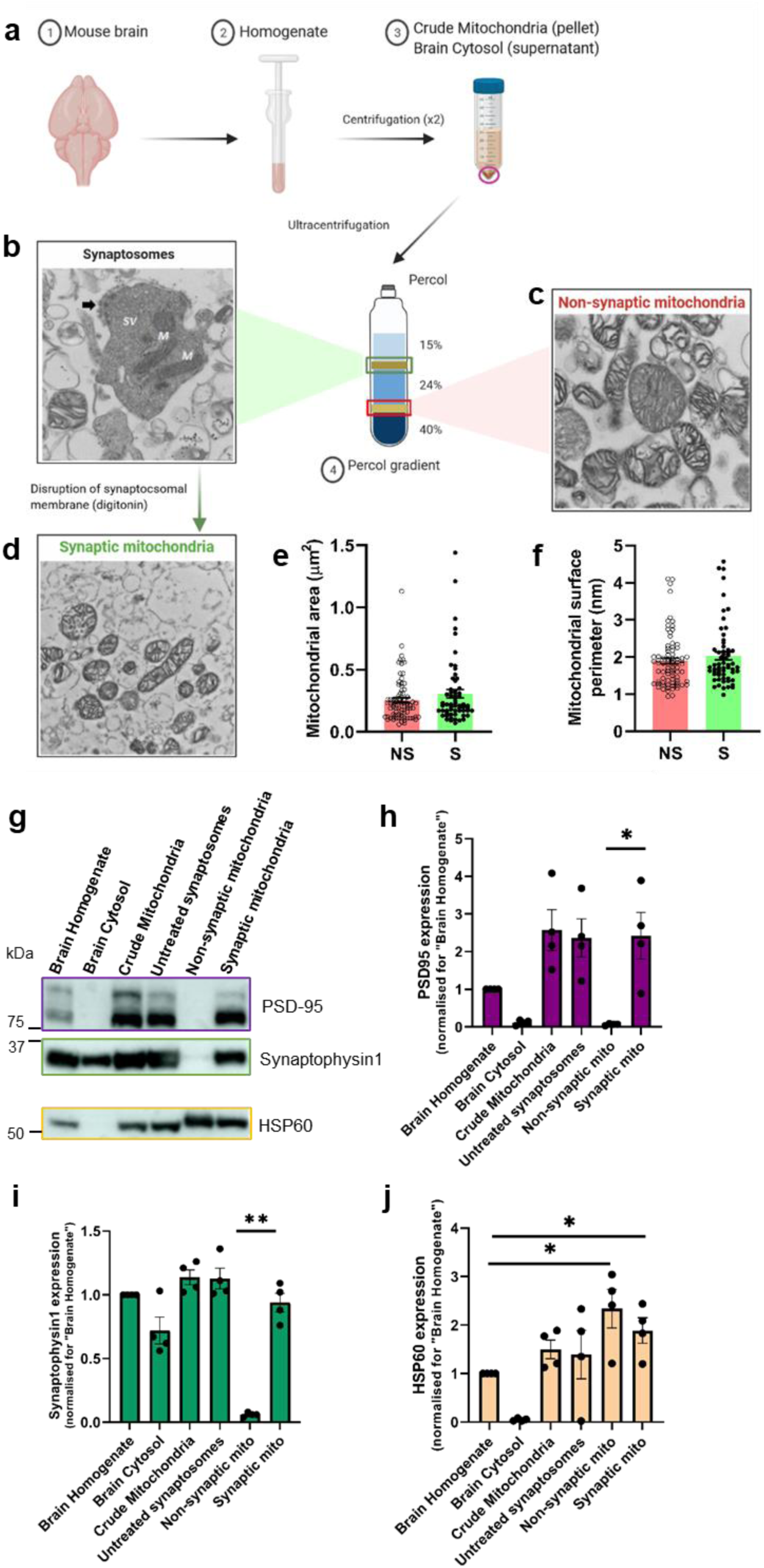
Isolation and morphological comparison of mouse synaptic and non-synaptic mitochondria. **a**, scheme of isolation procedure and representative TEM images of **b**, freshly isolated, undisrupted synaptosomes with synaptic vesicles (SV), mitochondria (M) and post-synaptic density (black arrow); **c**, non-synaptic mitochondria and **d**, synaptic mitochondria. (scale bar = 500 nm). Quantification of mitochondrial area (µm^2^) (**e**) and mitochondrial perimeter (nm) (**f**) from TEM images of non-synaptic and synaptic mitochondria (representative of 4 images per condition, from n=1 biological replicate). Expression (**g**) and quantification of post-synaptic PSD-95 (**h**), pre-synaptic Synaptophysin1 (**i**) and mitochondrial marker HSP60 (**j**) (n= 4 biological replicates). **e**, **f**, **h**, **i** and **j** show means ± S.E.M., determined by two-tailed Mann-Whitney test (**b**), two-tailed unpaired t-test (**b**) and two-tailed unpaired t-test with Welch’s correction (**h** - **j**), \**P* < 0.05, \*\**P* < 0.01.

Transmission electron microscopy (TEM) was used to assess the overall mitochondrial morphology (Fig. 1b-d; Extended Data Fig. 1b). In the untreated synaptosomal fraction (Fig. 1b; Extended Data Fig. 1b), we observe mitochondria (M), synaptic vesicles (SV) and post-synaptic densities (black arrow). Interestingly, several synaptosomes had one or more mitochondria, supporting the enrichment of synaptic mitochondria in these structures^18,24^. No significant differences were observed between synaptic (S) and non-synaptic (NS) mitochondria (Fig. 1e,f), challenging previously established ideas that synaptic mitochondria are smaller and rounder than non-synaptic mitochondria^17^. Synaptic mitochondria presented increased heterogeneity for area (0.25 ±0.020 for NS vs 0.31±0.034 for S; mean±SEM) and perimeter (1.90±0.079 for NS vs 2.03±0.109; mean±SEM).

Expression of mitochondrial protein HSP60 and pre- and post-synaptic markers Synaptophysin1 and PSD95, respectively, were assessed in the different fractions (Fig. 1g-j). As expected, the mitochondrial marker HSP60 was enriched in non-synaptic and synaptic mitochondria in comparison to the brain homogenate (Fig. 1g,j). The synaptic markers, PSD95 and Synaptophysin1 were close to absent in non-synaptic mitochondria, whereas they were detected in synaptic mitochondria, validating the enrichment of each mitochondrial fraction (Fig. 1g-i). The astrocyte marker GFAP, was not enriched in the mitochondrial fractions further validating the enrichments (Extended Data Fig. 1a).

To demonstrate that digitonin treatment applied on the synaptosomal fraction maintained the mitochondria membrane integrity^25^, lysed synaptosomal fraction were collected (hereafter, synaptosomal membrane) and probed for Citrate Synthase (CS), a mitochondrial matrix protein, VDAC1 and Mitofusin1 (MFN1), both outer mitochondrial membrane proteins, and SNAP25, a non-vesicular synaptosomal protein (Extended Data Fig. 1c-f). CS, VDAC1 and MFN1 are significantly enriched in synaptic mitochondria but not in synaptosomal membrane fractions, whereas SNAP25 is present in both, but increased in synaptosomal membranes. These results indicate that, at this concentration of digitonin, we obtained an enriched fraction of morphological intact synaptic mitochondria.

To elucidate the unique properties of mitochondria that reside at the synapse in comparison to non-synaptic mitochondria, a bioenergetic and biochemical profile was established from both fractions.

### Synaptic mitochondria present higher bioenergetic flexibility

The bioenergetics profile of synaptic and non-synaptic mitochondria was assessed by tracing the oxygen consumption rate (OCR)^26^. Basal respiration shows that both preparations are metabolically active with non-synaptic mitochondria respiring more than synaptic mitochondria (1206 ± 56,4 pmol/min versus 728,3 ± 37,7 pmol/min, *P*<0,0001) (Fig. 2a-b). To guarantee that ADP was not rate limiting and to boost coupled respirations^26^, we applied this substrate in the measurements. The resulting increase in respiration was significantly higher in synaptic mitochondria compared to non-synaptic mitochondria (64,6% ± 4,9% versus 32,9% ± 4,4%, *P*<0,0001) (Fig. 2d,e). When complex V is blocked with oligomycin, the effect was higher in synaptic mitochondria (51,1% ± 1,1% versus 31,1% ± 3,5%, *P*<0,0001) (Fig. 2f) suggesting higher flexibility of synaptic mitochondria to produce ATP. The increase in maximal uncoupler-stimulated respiration was also significantly higher in synaptic mitochondria than in non-synaptic mitochondria (67,7% ± 5,8% versus 17,0% ± 3,4%, *P*<0,0001) (Fig. 2g). After electron transport chain block with Antimycin A, both the synaptic and non-synaptic mitochondrial OCR became drastically decreased to a similar level indicating that none of the mitochondrial fractions have a significant source of non-mitochondrial respiration (Fig. 2c).

**Fig. 2.**
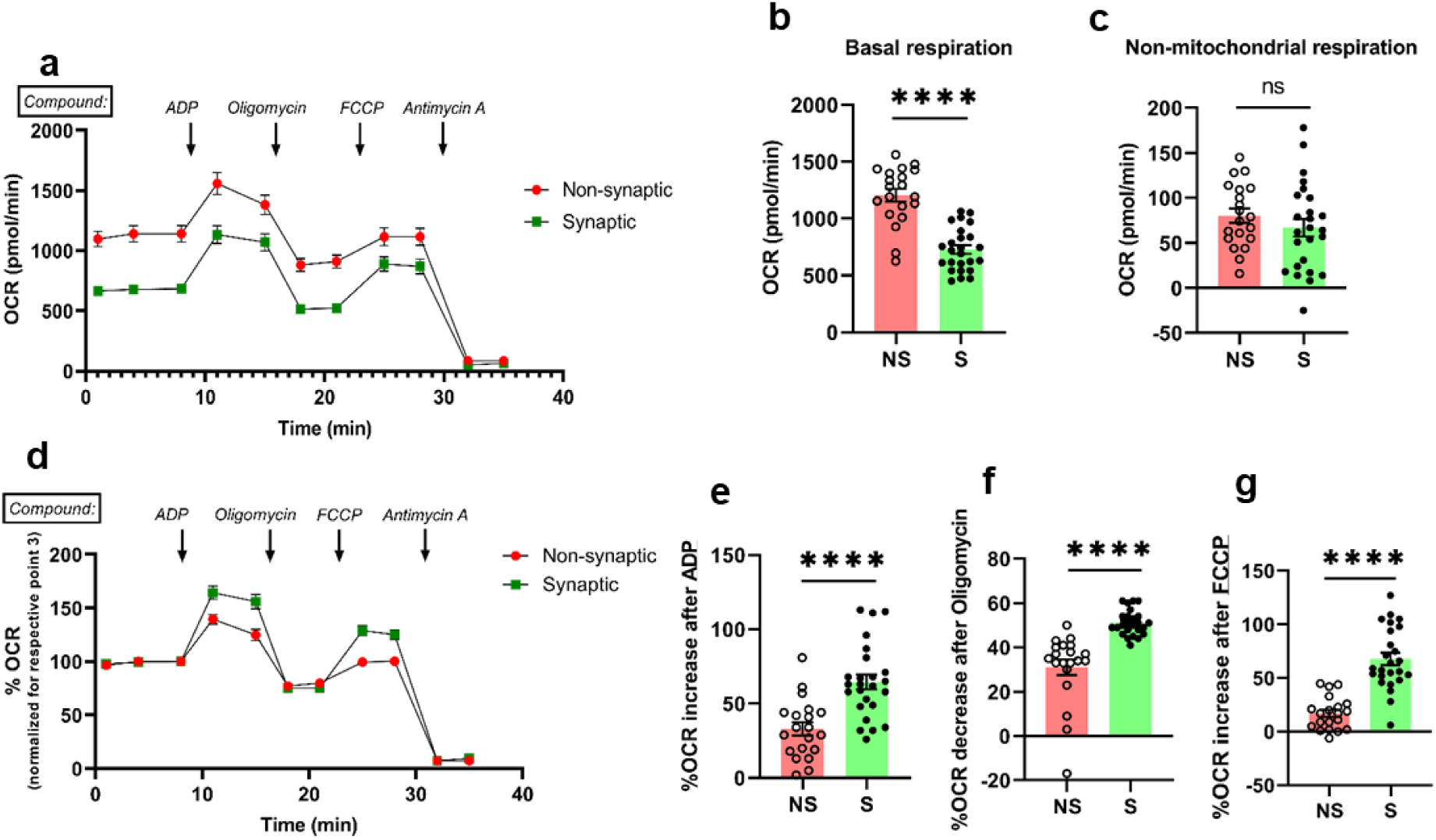
Freshly isolated brain mitochondria are bioenergetically active. **a** and **d**, oxygen consumption rate (OCR) curves and quantification of 8ug non-synaptic and synaptic mitochondria showing basal conditions (**b**) and non-mitochondrial respiration (**c**). Percentage of OCR increase after ADP (**e**) and FCCP (**g**) and OCR decrease after Oligomycin (**f**) (n=6). **a** - **g** show means ± S.E.M., determined by two-tailed unpaired t-test (**b**, **c** and **e**), and two-tailed unpaired t-test with Welch’s correction (**f**, **g**), \**P* < 0.05, \*\*\*\**P* < 0.0001.

Thus, while synaptic mitochondria present a reduced basal respiration, they react at a higher magnitude upon different respiratory stimuli. This supports the hypothesis that mitochondria at synapses are in a “low-oxidative phosphorylation” state in basal synaptic transmission, but upon high-demanding synaptic transmission^11,12^ are capable to adapt and contribute to synaptic activity.

### Synaptic mitochondria have increased OxPHOS complex activity, except for Complex I

We assessed the enzymatic activities of the individual respiratory chain complexes (RCCs) I, II, III, IV and V (Fig. 3a-e), as well as, the combined activity of Complex I+III and Complex II+III (Fig. 3f,g) of synaptic and non-synaptic mitochondria. Interestingly, the activities of Complex IV and, more drastically, of Complex V were increased in synaptic mitochondria (Fig. 3d,e). However, Complex I activity (Fig. 3a) was decreased in synaptic mitochondria, while no significant changes for Complex II (Fig. 3b) and Complex III (Fig. 3c) were observed, pointing to a differential activity of the RCCs in synaptic mitochondria^14,28^. Although no significant change of Complex II+III was observed in synaptic mitochondria (Fig. 3g), a significant increase in the enzymatic activity of combined Complex I+III was evident in synaptic mitochondria (Fig. 3f).

**Fig. 3.**
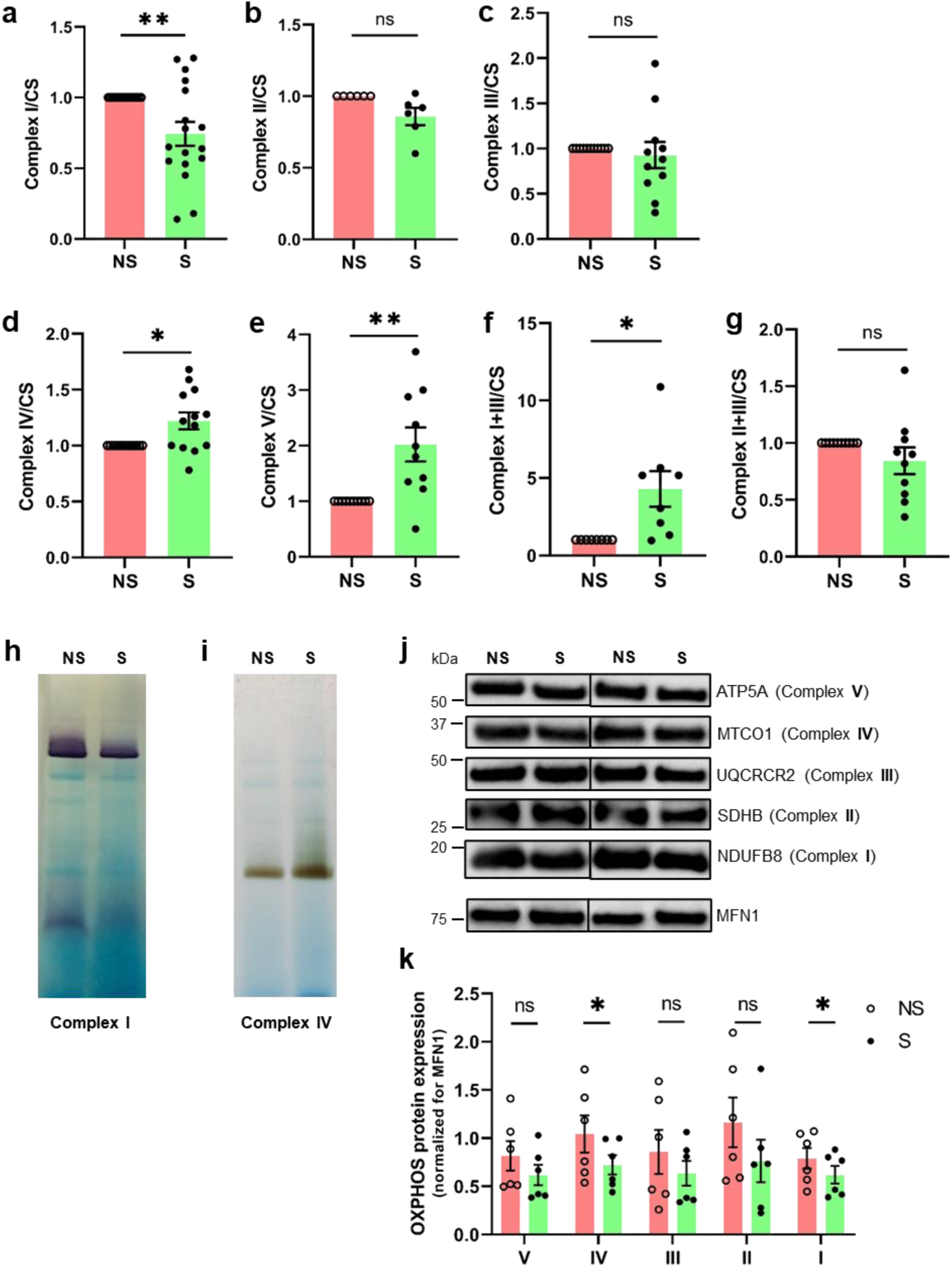
Synaptic mitochondria show different individual and combined activity of the respiratory chain complexes. Enzymatic activities of the individual respiratory chain complexes I, II, III, IV and V (**a** - **e**), as well as, combined activity of Complex I+III (**f**) and Complex II+III (**g**) were assessed in synaptic mitochondria and normalized for the respective activity of non-synaptic mitochondria. All activities were normalized to Citrate Synthase (CS) activity (n=6-17). In-gel activity of Complex I in DDM-treated brain mitochondria for Complex I (**h**) and Complex IV enzymatic activity (**i**) in synaptic mitochondria (S) in comparison to non-synaptic mitochondria (NS) (n=4). Protein expression (**j**) and quantification (**k**) of a subunit of each mitochondria RCCs in synaptic (S) and non-synaptic (NS) mitochondria. Bands intensity were normalized for Mitofusin1 (MFN1) expression (n=6). **a-g**, **k** show means ± S.E.M., determined by two-tailed unpaired t-test with Welch’s correction (**a-g**) and two-tailed paired t-test (**k**). \**P* < 0.05, \*\**P* < 0.01.

The decrease of individual Complex I activity in synaptic mitochondria was confirmed by in-gel activity assay (Fig. 3h), where DMM treated mitochondria were used (condition that only allows analysis of individual RCCs)^29^, whereas Complex IV activity was increased in synaptic mitochondria (Fig. 3i). Additionally, expression of a subunit from each mitochondrial RCCs was assessed with antibodies against ATP5A (V), MTCO1 (IV), UQCRCR2 (III), SDHB (II) and NDUFB8 (I). Both Complex IV and I subunits protein levels in synaptic mitochondria (Fig. 3j,k).

Collectively, these data suggests that synaptic mitochondria have a specific and distinct bioenergetic profile than other brain mitochondria, with a higher activity of Complex IV, V and I+III but decreased enzymatic activity of Complex I.

### Differential Complex I levels and function in synaptic mitochondria

Mammalian Complex I is the largest mitochondrial respiratory chain complex and contains several redox centers where electrons from NADH are accepted by ubiquinone and NADH is oxidized to NAD^+^. Functionally, Complex I can be divided in 3 modules: N-module, where NADH is oxidized and electrons are accepted to FMN; Q-module, where the ubiquinone Coenzyme Q10 is reduced to deliver electrons to Complex III; P-module, where protons are pumped to the mitochondrial intermembrane space, contributing to the build-up of the mitochondrial membrane potential (Fig. 4a).

**Fig. 4.**
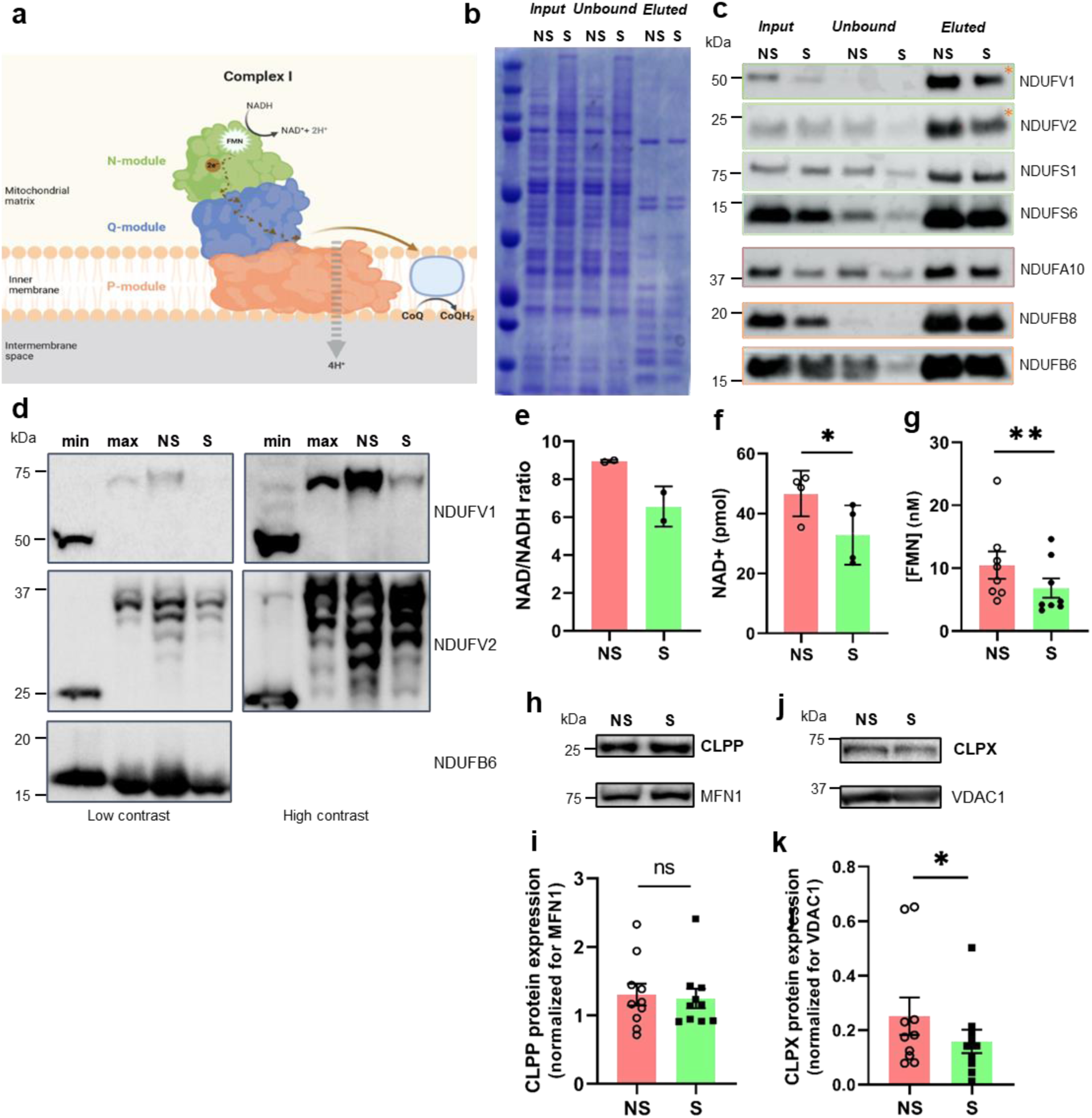
Differential Complex I levels and function in synaptic mitochondria. Schematic (**a**) of Complex I structural composition: N-module (green), where NADH is oxidized to NAD^+^ and flavin mononucleotide (FMN) is bound; Q-module (blue), where electrons are transfered to ubiquinone; P-module (orange) localised in the mitochondrial inner membrane and where protons (H^+^) and pumped to the mitochondrial inner membrane space. Coomassie blue staining (**b**) and immunoblot (**c**) of subunits from different modules of Complex I in co-immunoprecipitation fractions from synaptic (S) and non-synaptic (NS) mitochondria (n=2). (**d**) Inverse Redox shift assay followed by immunoblot of NDUFV1, NDUFV2 and NDUFB6 in synaptic (S) and non-synaptic (NS) mitochondrial fractions (n=4). NAD/NADH ratio (n=2) (**e**), NAD+ quantification (n=4) (**f**) and FMN content determination (n=8) (**g**) of synaptic and non-synaptic mitochondrial-enriched fractions. CLPP expression (**h**) and quantification (**i**) (n=9) and of CLPX (**j** - **k**) (n=10) in synaptic and non-synaptic mitochondrial fractions. **e, f, g, i** and **k** show means ± S.E.M., determined by two-tailed Wilcoxon test (**g** and **k**) and two-tailed paired t-test (**f**, **i**), \**P* < 0.05, \*\**P* < 0.01.

Complex I is also one of the major producers of mitochondrial reactive oxygen species (ROS)^30^.

We immunoprecipitated Complex I from synaptic and non-synaptic mitochondria and compared the expression of Complex I subunits from the 3 different modules: NDUFV1, NDUFV2, NDUFS1 and NDUFS6 (N-module subunits); NDUFA10 (Q/P-module interface) and NDUFB8 and NDUFB6 (P-module subunits) (Fig. 4b-c). In the “input” fractions, there is an overall decrease in protein levels of Complex I subunits in the synaptic mitochondria (Fig. 4c). In the “eluted” samples decrease in protein levels was mainly observed for the two core subunits of the N-module, NDUFV1 and NDUFV2 (Fig. 4c, *).

To assess whether decreased protein levels was due to protein instability, we next analysed the tertiary conformation of these subunits by assessing cysteine modifications using inverse redox shift assays^31^. Mitochondrial samples are first exposed to N-ethylmaleimide (NEM) which binds to accessible cysteine residues and is represented by a lower molecular weight band similar to minimum shift control lane (“min” lane). On the other hand, proteins with inaccessible cysteine will present high molecular weight bands similar to maximum shift control lane (“max” lane). In Figure 4d, lower molecular weight bands were not observed for NDUFV1 in both non-synaptic (NS) and synaptic (S) mitochondria. For the NDUFV2 subunit, intermediate molecular weight bands were observed in both maximum, non-synaptic and synaptic mitochondria. Additionally, no exclusive band was shown in synaptic mitochondria in comparison with non-synaptic mitochondria. As expected, no shift bands were observed between minimum, maximum and steady-state samples for NDUFB6, a protein that lacks cysteine residues. As Complex I inactivation can be derived from irreversible conformational changes of N-module subunits^32^, our results indicate that this is not the case in synaptic mitochondria. Commonly, Complex I deficiencies are accompanied by a lower NAD^+^/NADH ratio^33^. Assessment of the NAD^+^/NADH ratio in non-synaptic and synaptic mitochondria (Fig. 4e) indicated that synaptic mitochondria have a tendency for lower NAD+/NADH ratio and a significantly decreased NAD+ concentration (Fig. 4f). Additionally, the amount of FMN in intact mitochondria was determined and our data shows that synaptic mitochondria indeed have lower FMN content than non-synaptic mitochondria (Fig. 4g), suggesting that lower Complex I activity (Fig. 3a,h) is accompanied by lower function and expression of N-module subunits.

Complex I N-module turnover occurs at a higher rate when compared to other Complex I modules^31,34^. This turnover depends on the mitochondrial ClpXP protease complex, which recognizes and degrades impaired core N-module proteins^31,35^. Upon absence of CLpXP complex, N-module turnover is diminished and leads to the accumulation of inactive CI subunits which interfere with the adequate assembly and activity of Complex I^31^. ClpPX consists of a CLPP and a CLPX component. CLPP was not different between NS and S mitochondrial (Fig. 4h,i), however, CLPX protein levels were decreased in synaptic mitochondria (1.85 fold) (Fig.4j,k). These lower levels of CLPX can contribute to a “stalling” mechanism of N-module turnover in synaptic mitochondria. However, these lower levels of CLPX do not seem to lead to an accumulation of N-module subcomplexes (Extended Fig. 2a), as observed in CLPP-deficient mouse hearts^31^. NDUFAF2 has been proposed to be an assembly factor that accumulates in pre-assembled Complex I (pre-CI) when N-module incorporation is impaired^36,37^. If pre-CI were to accumulate in synaptic mitochondria, an increased NDUFAF2 expression would be expected. However, synaptic mitochondria fractions showed a decreased NDUFAF2 expression when compared to non-synaptic mitochondria (Extended Data Fig. 4b,c). Interestingly, in NDUFAF2 KO cells, Complex I can be fully assembled but its function seems to be impaired^38^.

At this point, we were able to unravel a N-module expression and function signature in synaptic mitochondria which could be causing the decreased Complex I enzymatic function in synaptic mitochondria and may constitute a protective mechanism that aims to maintain Complex I-produced ROS reduced under basal synaptic activity, as that it has been observed that, upon synaptic stimulation, NDUFV1 expression is enhanced at synapse^21^.

### Integration of Complex I in supercomplexes is increased in synaptic mitochondria

RCCs can be arranged in closer proximity forming supercomplexes, that overall contribute to a lower electron diffusion distance between complexes ^39,40^, leading to an increased ATP production efficiency with lower ROS production^41,42^. Our data reveal reduced individual Complex I activity (Fig. 3a) and increased Complex I+III combined enzymatic activity (Fig. 3b) in synaptic mitochondria. To assess Complex I distribution in individual or supercomplexes, we treated mitochondrial-enriched fractions with digitonin, a weak nonionic detergent that maintains supercomplex formation^29^. Synaptic mitochondria present decreased levels of individual Complex I but similar levels of supercomplex integrated-Complex I when compared to non-synaptic mitochondria (Fig. 5a-d). Calculating the ratio of Complex I distribution assessing core subunits of N-module (Fig. 4a), NDUFV1 (Fig. 5b,g) and NDUFV2 (Fig. 5c,h), an accessory subunit of N-module, NDUFS6 (Fig. 5a,f) and a subunit of P-module (Fig. 4a), NDUFB6 (Fig. 5d,i), synaptic mitochondria present increased supercomplex integrated-Complex I. No differences in Complex II expression were observed (Fig. 5e). Second-dimension BN/SDS-PAGE (Fig. 5j) of digitonin-solubilized synaptic and non-synaptic mitochondria, confirmed NDUFV1 subunit in supercomplexes and individual Complex I in non-synaptic mitochondria, while NDUFV1 was only detected in supercomplexes of synaptic mitochondria. The Complex III subunit, UQCRFS1, was present in CIII_2_ and supercomplex structures in both mitochondrial populations.

**Fig. 5.**
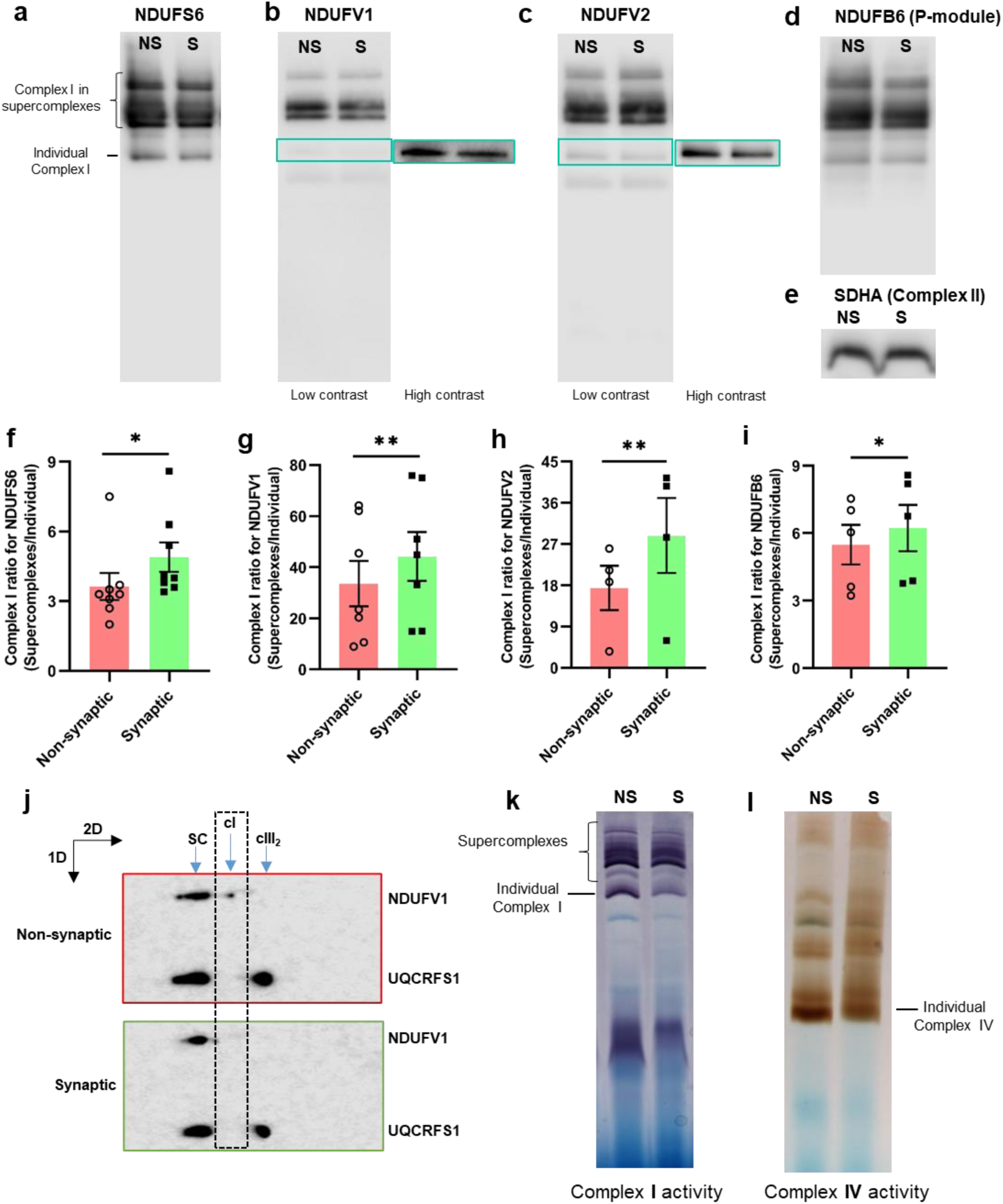
Synaptic mitochondria show higher Complex I integration in supercomplexes than non-synaptic mitochondria. Immunodetection of Complex I subunits in BN-PAGE of digitonin-treated synaptic (S) and non-synaptic (NS) mitochondria fractions: NDUFS6 (**a**), NDUFV1 (**b**), NDUFV2 (**c**); NDUFB6 (**d**); and Complex II subunit SDHA (n=3) (**e**). Ratio quantification of Complex I integration in supercomplexes to individual Complex I of NDUFS6 (n=8) (**f**), NDUFV1 (n=7) (**g**), NDUFV2 (n=4) (**h**); NDUFB6 (n=5) (**i**). (**j**) Second-dimension BN/SDS-PAGE of digitonin-solubilized synaptic and non-synaptic mitochondria, followed by immunodetection of the NDUFV1 (Complex I) and UQCRFS1 (Complex III subunit). In-gel enzymatic activity of Complex I (**k**) and Complex IV (**l**) in digitonin-treated synaptic (S) and non-synaptic mitochondria (NS) (n=4). **f-i** show means ± S.E.M., determined by two-tailed paired t-test, \**P* < 0.05, \*\**P* < 0.01.

In-gel activity assays revealed a reduction in individual Complex I activity in synaptic mitochondria, while supercomplex presented no significant difference between the two mitochondrial pools (Fig. 5k). A similar trend is observed for Complex IV activity (Fig. 5l). These data indicates that, although overall Complex I protein levels and individual activity is decreased in synaptic mitochondria, the existent Complex I seems to be almost totally organized into supercomplexes. This feature is a plausible justification for a higher response capacity to energetic demands, previously observed in synaptic mitochondria (Fig. 2d-g).

### Synaptic mitochondria show differential arrangement of Complex V

Besides differences in Complex I activity, Complex V presented a higher enzymatic activity in synaptic mitochondria when compared to non-synaptic mitochondria (Fig. 3e). Complex V can be organized as monomers, dimers and oligomers^43–45^ and these structures improve overall ATP synthesis capacity^45^. To decipher Complex V organization in synaptic mitochondria, BN-PAGE/WB analyses of mitochondria subjected to digitonin/protein ratios of 10:1 and 2:1 (Fig. 6a) were performed. As expected, dimeric and oligomeric forms of Complex V were maintained in lower digitonin/protein (2:1) ratio^46^. Synaptic mitochondria present an increased incorporation of Complex V in oligomers than non-synaptic mitochondria (Fig. 6a,b), and a subtle tendency for increased dimeric forms also occurs in synaptic mitochondria (Fig. 6a,c), suggesting that the supramolecular organization of Complex V in dimers and oligomers is favoured in synaptic mitochondria, which corroborate higher enzymatic activity of Complex V in synaptic mitochondria (Fig. 3e).

**Fig. 6.**
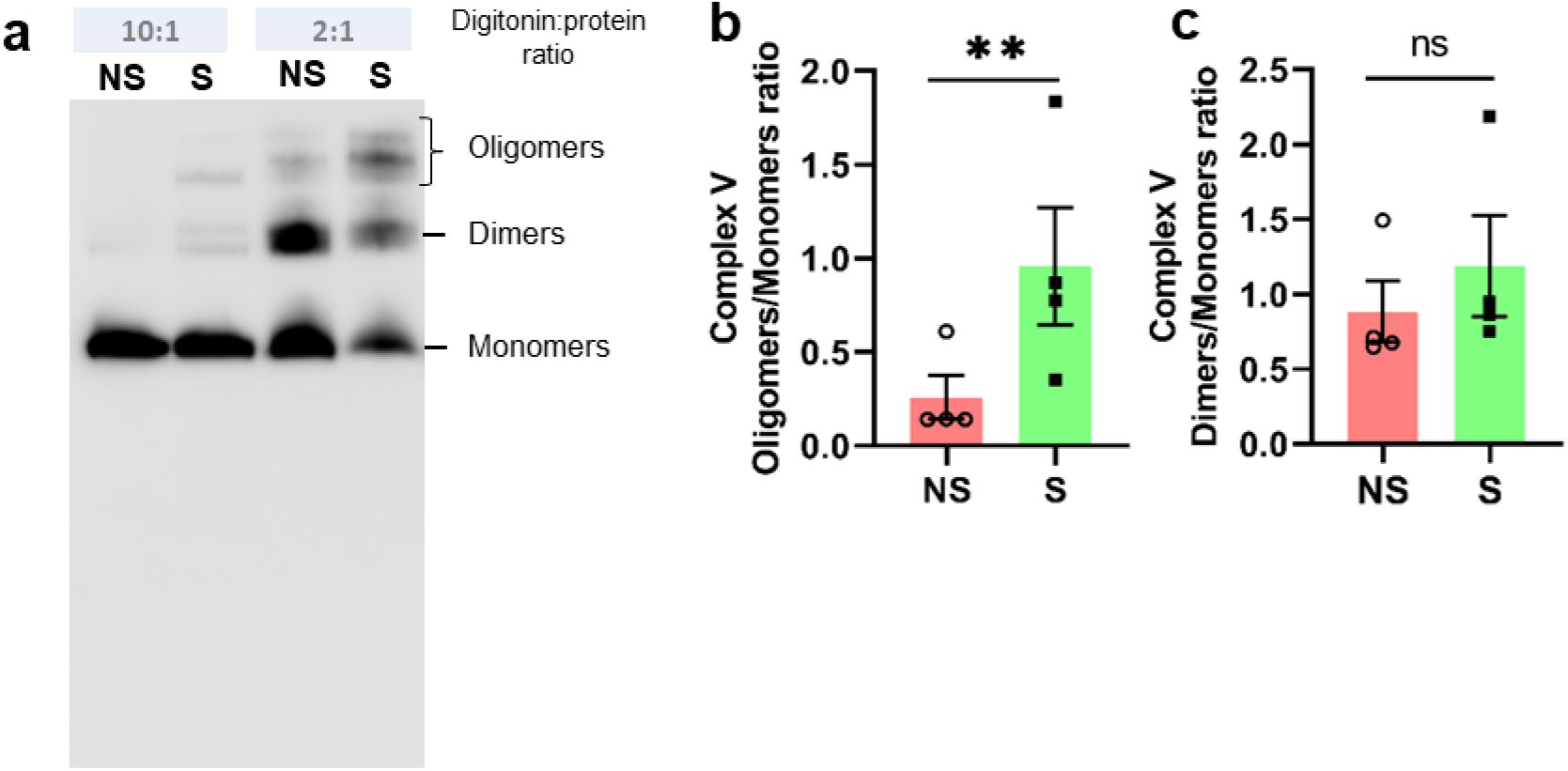
Synaptic mitochondria show differential arrangement of Complex V. BN-PAGE (**a**) of digitonin-solubilized synaptic (S) and non-synaptic (NS) mitochondria fractions, using different digitonin:protein ratios (10:1 and 2:1) followed by immunodetection of Complex V subunit (ATP5B) (n=3). Ratio quantification of Complex V organization in oligomers to monomers (**b**) and dimers to monomers (**c**). **b** and **c** show means ± S.E.M., determined by two-tailed paired t-test, \**P* < 0.05, \*\**P* < 0.01.

In sum, a preferential Complex I incorporation in supercomplexes and an increase in Complex V oligomeric organization may contribute to the increased bioenergetic flexibility of synaptic mitochondria.

Overall, mitochondria need to deal with a wide and dynamic range of metabolic conditions and synaptic mitochondria are a clear example: they have to be able to adapt to resting and high stimulation environments which translate into differential energetic needs overtime. Thorough analysis of synaptic mitochondria has revealed that even though a lower respiration rate is observed in resting conditions, upon stimulation of the respiratory chain a higher energetic flexibility is observed. Interestingly, even though a reduced Complex I enzymatic activity, accompanied by a lower FMN and NAD+ content was observed, synaptic mitochondria present increased Complex I integrated supercomplexes clearly supporting an intrinsic bioenergetic-favourable signature of synaptic mitochondria.

Upon intense stimulation conditions, where a maximum energetic output is necessary (mainly through OxPHOS^11^), more N-module and full Complex I would be generated to sustain those high energetic demands, which is supported by recent study showing that *ex vivo* glutamatergic stimulation of synaptosomes leads to increased NDUFV1 expression^21^. Furthermore, in a Complex I deficiency model, where both individual and supercomplexes-integrated Complex I were reduced, it was demonstrated that the early brain energy deficits were detected in synaptosomes whereas non-synaptic mitochondria remained unaffected^47^, supporting a critical role of Complex I in synaptic energy metabolism.

The preferential integration of Complex I into supercomplexes was previously detected when comparing neuronal and astrocytic mitochondria^42^. Our data not only corroborates these findings but further indicates that this feature may actually be a signature of mitochondria at synapse, supporting the concept that mitochondria inside a cell are functionally distinct from mitochondria from other subcellular compartments – the so called “mitochondrial microheterogeneity”^23,48^.

As the onset of neurodegenerative disorders is associated with synaptic energy deficits, mitochondrial dysfunction and ultimately synaptic degeneration prior to neuronal loss^49–51^, the definition of the bioenergetics profile of synaptic mitochondria under physiological conditions could set the ground to understand the energy deficits that occur in neurodegenerative disorders, even in early asymptomatic stages.

## ACKNOWLEDGEMENTS

The authors are grateful to all VMorais lab members for their input and technical assistance, especially to Carlos Cardanho-Ramos and Bernardo Antunes. The authors are grateful to the Animal and Comparative Pathology (especially Tânia Carvalho and Andreia Pinto) facilities, and we also acknowledge the funding PPBI-POCI-01-3700145-FEDER-022122. The authors are also thankful to LLopes and EGomes labs at GIMM for their help in provision of antibodies and animals.

## AUTHOR CONTRIBUTIONS

Conceptualization – A.F.-P.; V.A.M; Methodology – A.F.-P.; K. C.; Investigation – A.F.-P.; Formal analysis: A.F.-P.; Data curation – A.F.-P.; Writing – A.F.-P.; V.A.M.; Supervision – V.A.M.; Funding acquisition – V.A.M.

## MATERIAL AND METHODS

### Isolation of mouse brain mitochondria

All the animal procedures were approved by the institutional ethics committee - Animal Welfare Body of GIMM (ORBEA-GIMM), and authorized and licensed by DGAV - Direcção Geral de Alimentação e Veterinária, the Portuguese Competent Authority for Animal Health. The study was carried out in compliance with the ARRIVE guidelines^52^.Brains were isolated from 8 weeks old C57BL/6J female mice and homogenized with a rotating Teflon 30 ml Potter-Elvehjem (15-20 strokes, 800 rpm) in a cold isolation buffer (IB: 10mM HEPES, 225mM sucrose, 75mM D-Mannitol and 1mM EGTA, pH7.4) followed by centrifugation at 600 xg, 10min, 4°C. Brain homogenate fraction was collected and further centrifuged at 15,000 xg, 10min, 4°C. Pellet (crude mitochondrial fraction) was ressuspended in 15% Percoll and applied on a 40% and 24% Percoll cushion and centrifuged in a fixed rotor at 31,000 xg, 9min, 4°C, with maximum acceleration and slow deceleration. Top white phase (mostly myelin) was removed and synaptosomes (in the interphase 15-24% Percoll) and non-synaptic mitochondria (in the interphase 24%-40% Percoll) were collected. In order to collect the synaptic mitochondria, synaptossomes were incubated with 0,6mg/ml digitonin (adapted from ^25^). Protein concentration was determined using Pierce™ BCA Protein Assay as described by the manufacturer (Thermo Fisher Scientific, 23225).

### Oxygen consumption rate on isolated brain mitochondria

Mitochondrial function was evaluated by measuring the oxygen consumption rate (OCR) in the XF24 Extracellular Flux Analyzer (Seahorse Bioscience, Agilent). Freshly collected synaptic and non-synaptic mitochondrial fraction (8µg of total protein concentration) were ressuspended in mitochondrial assay solution (MAS) ^26^ and plated in a 24-well Seahorse plate followed by centrifugation 2,000 xg, 2min, 4°C. Three measurements were collected for basal respiration, followed by two measurements after addition of 2mM ADP, followed by two measurements after addition of 20μM oligomycin, followed by two measurements after addition of 20μM Carbonyl cyanide-4- (trifluoromethoxy) phenylhydrazone (FCCP), followed by two measurements after addition of 20μM antimycin A.

### Respiratory complexes enzymatic activity assays

For this, 2 to 50μg of protein from synaptic and non-synaptic mitochondria was used. Individual Complex I (NADH:decylubiquinone oxidoreductase, rotenone sensitive, Complex II (succinate: decylubiquinone DCPIP reductase, malonate sensitive), Complex III (decylubiquinol:cytochrome c oxidoreductase, antimycin A sensitive), Complex IV (cytochrome c oxidase), Complex V (ATPase, oligomycin sensitive), Complexes I+III (NADH:cytochrome c oxidoreductase, rotenone and antimycin A sensitive), Complexes II+III (succinate:cytochrome c reductase activity) and citrate synthase enzymatic activity were measured as described ^53,54^. Values were plotted according to the ratio between the specific enzymatic activity of each complex activity and normalized to citrate synthase enzymatic activity.

### In-gel activity of respiratory chain complexes I and IV

250µg of mitochondria were ressuspended in 750mM aminocaproic acid, 50mM Bis-Tris/HCl pH7.0, 20µM PMSF and 1% DDM or Digitonin, incubated on ice for 30min (DDM) or 1h (Digitonin), centrifuged at 100,000 xg for 15min and supernatants were collected and protein concentration was determined. Next, 0.25% Coomassie G-250 ressuspended in 750mM aminocaproic acid was added to 30µg of each mitochondrial fraction, similar to ^54^.

### Immunoprecipitation of Complex I subunits

Complex I was immunocaptured from brain mitochondrial enriched fractions as described in ^55^. Samples were further analyzed by immunoblotting.

### Immunoblot and BN-PAGE analysis

For immunoblot analysis, proteins were separated by SDS-PAGE and electrophoretically transferred onto 0.2µm nitrocellulose membranes. Incubation with primary antibodies was performed overnight at 4°C using dilutions listed in Table 1. The following secondary antibodies were used: Horseradish peroxidase (HRP) conjugated anti-rabbit and anti-mouse (Bio-Rad) at 1:10,000. Detection was done using the chemiluminescent ECL-Plus detection kit (Amersham) on a digital Amersham Imager 680 (GE Healthcare). Semi-quantification of band intensity was determined using Image Lite Studio 5.2 software.

**Table 1:**
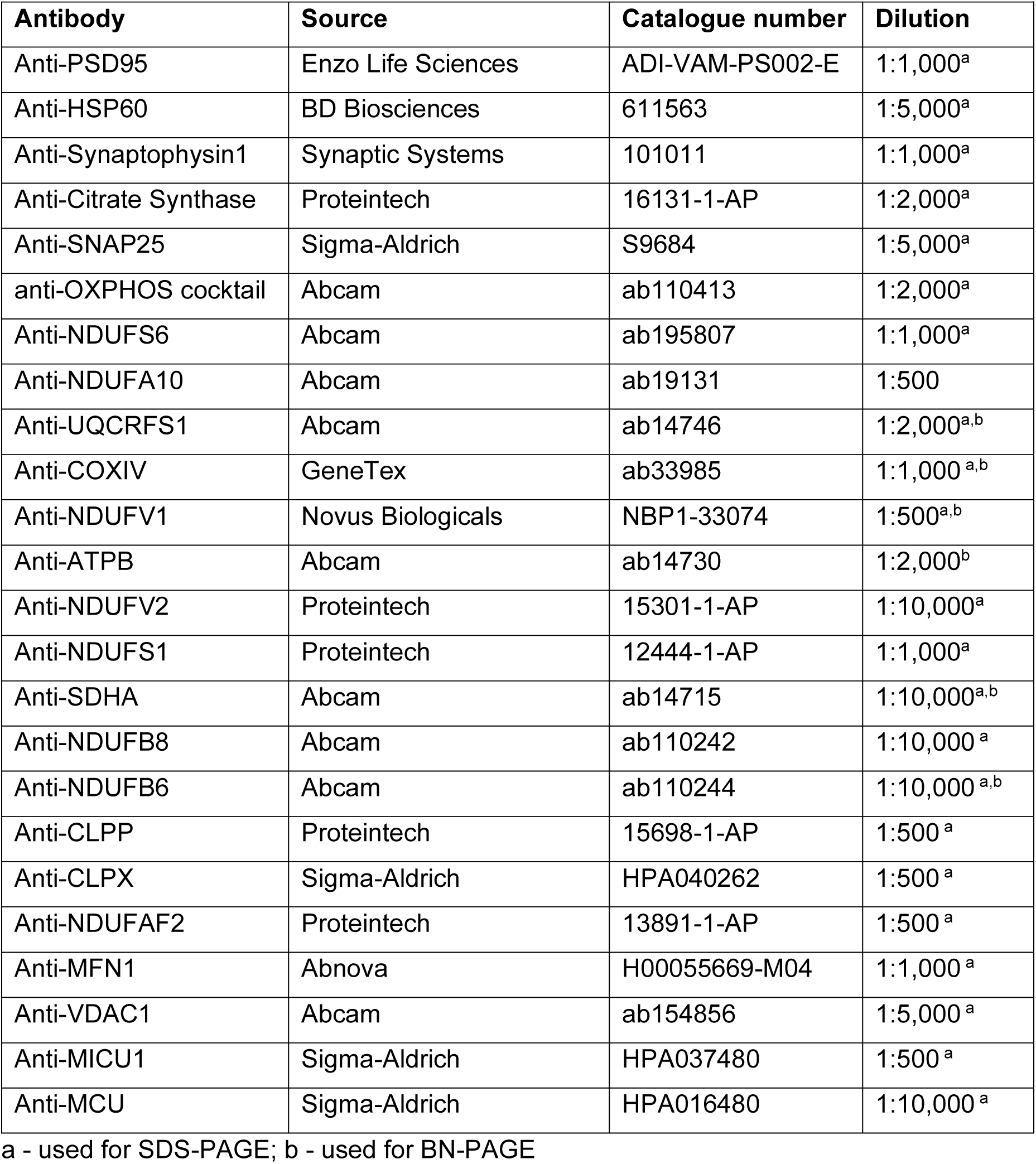
Primary Antibodies used in this study.

Samples for blue-native gel electrophoresis (BN-PAGE) were prepared from brain isolated mitochondria. For the solubilization, 10µg digitonin/µg protein was used. Approximately, 25µg of protein was loaded on a 3–12% Native PAGE Bis-Tris gels (Thermo Fisher Scientific) and stained with 0.1% Coomassie R-250 solution. Proteins were electrophoretically transferred onto 0.45µm PVDF membranesand immunodetection was performed as previously described using commercial antibodies (Table 1).

### NAD/NADH quantification

Using the colorimetric NAD/NADH Assay, 250µg freshly isolated brain mitochondrial enriched fractions were processed according to the manufacturer’s instructions (ab65348, Abcam). Absorbance at 450nm was measured during 1 to 5 hours. The concentration of total NADH, and NAD and NADH concentrations were obtained by comparison to a NADH standard curve. The values were plotted as the ratio between the calculated concentration of NAD+ and NADH.

### FMN content determination

1mg/ml of brain isolated mitochondria were mixed with an equal volume of 15% TCA for 10min on ice, centrifuged at 10,000 x g for 10min 4 °C and neutralized with 4M K_2_HPO_4_. Fluorescence was measured at excitation/emission 450/525nm. First, the fluorescence was recorded at acidic conditions, and then under neutral pH conditions (pH ∼ 7.4) obtained by addition of 20μl of 4M K_2_HPO_4_. Freshly prepared FMN standard solutions were used to calibrate the fluorescence signal (adapted from ^31,56^).

### Inverse Redox shift assay

Inverse Redox shift protocol was adapted from ^31^. Briefly,25µg of isolated mitochondria were used for each sample. For minimum shift and maximum shift sample, mitochondria were washed with PBS. For steady-state samples, mitochondria were washed and incubated with ice-cold 20mM N-ethylmaleimide (NEM) for 10min on ice. Incubated samples were concentrated using a Microcon-10kDa columns (Merck) and NEM-PBS was replaced with PBS. Concentrated proteins were ressuspended in ice-cold 8% TCA and incubated overnight at −20°C. Precipitated proteins were centrifuged for 15min at 20,000 xg, 4°C. Protein pellets were washed with ice-cold 5% TCA and solubilized in 1x Laemmli Sample buffer with 10mM Tris(2-carboxyethyl)phosphine (TCEP) with sonication. Samples were incubated for 15min at 45°C with vigorous agitation and cooled down at room temperature. Minimum shift samples were modified with 15mM NEM and maximum shift and steady-state samples were modified with 15mM mm(PEG)24. All samples were incubated at RT in the dark for 1h. Redox state was analyzed by SDS-PAGE immunoblotting.

### Electron microscopy

Mitochondrial suspensions were fixed in 2.5% glutaraldehyde in 0.1M sodium cacodylate buffer previously pre-warmed at room temperature for 1h followed by three washes in the same buffer. Post-fixation was performed with 1% (aq) osmium tetroxide for 1h on ice. En bloc staining was performed with 1% (aq) uranyl acetate for 30 min Samples were embedded into BEEM® capsules filled with EPOXI resin (Epon 812 from EMS) and baked for 48 hours at 68-70°C. Ultrathin sections (70nm) were obtained and collected into 1% formvar coated slot copper grids and post stained with 2% Uranyl acetate. Sections were examined in a Hitachi H7000 electron Microscope at a voltage of 80 Kv and the images were acquired using a Megaview III mid-mounted camera (Olympus) and post-processed using iTEM platform.

For analysis of perimeter and mitochondrial area, TEM images (.tiff format) were manually analysed based on the outer mitochondrial membrane (OMM) using Fiji/ImageJ software. The following inclusion criteria were applied: the entire mitochondrial 2D representation had to be within the image, not cutted in the limits and the OMM had to be clearly throughout the mitochondria. Note that in synaptic mitochondria’s TEM images the abundance of mitochondria seems to be lower than in non-synaptic mitochondria’s TEM images; however the samples of each fraction processed for TEM had not the same protein concentration and volume.

### Statistical analysis

Statistical analysis was performed using GraphPad Prism8.0 (GraphPad Software, Inc.). Each set of data values were subjected to normality tests (Shapiro-Wilk test and Kolmogorov-Smirnov test and assessment of Q-Q plots). When data groups assumed a gaussian (normal) distribution, parametric tests were applied, otherwise non-parametric tests were used. Statistical details of experiments, including the statistical tests used and the number of biological replicates (n), are noted in figure legends. Data are presented as mean ± SEM. Differences were considered significant if p-value was lower than 0.05.

