## Extended Data for "Bioenergetic signature of Synaptic mitochondria"

**TITLE**:

**CORRESPONDING AUTHOR**

* Vanessa A. Morais,

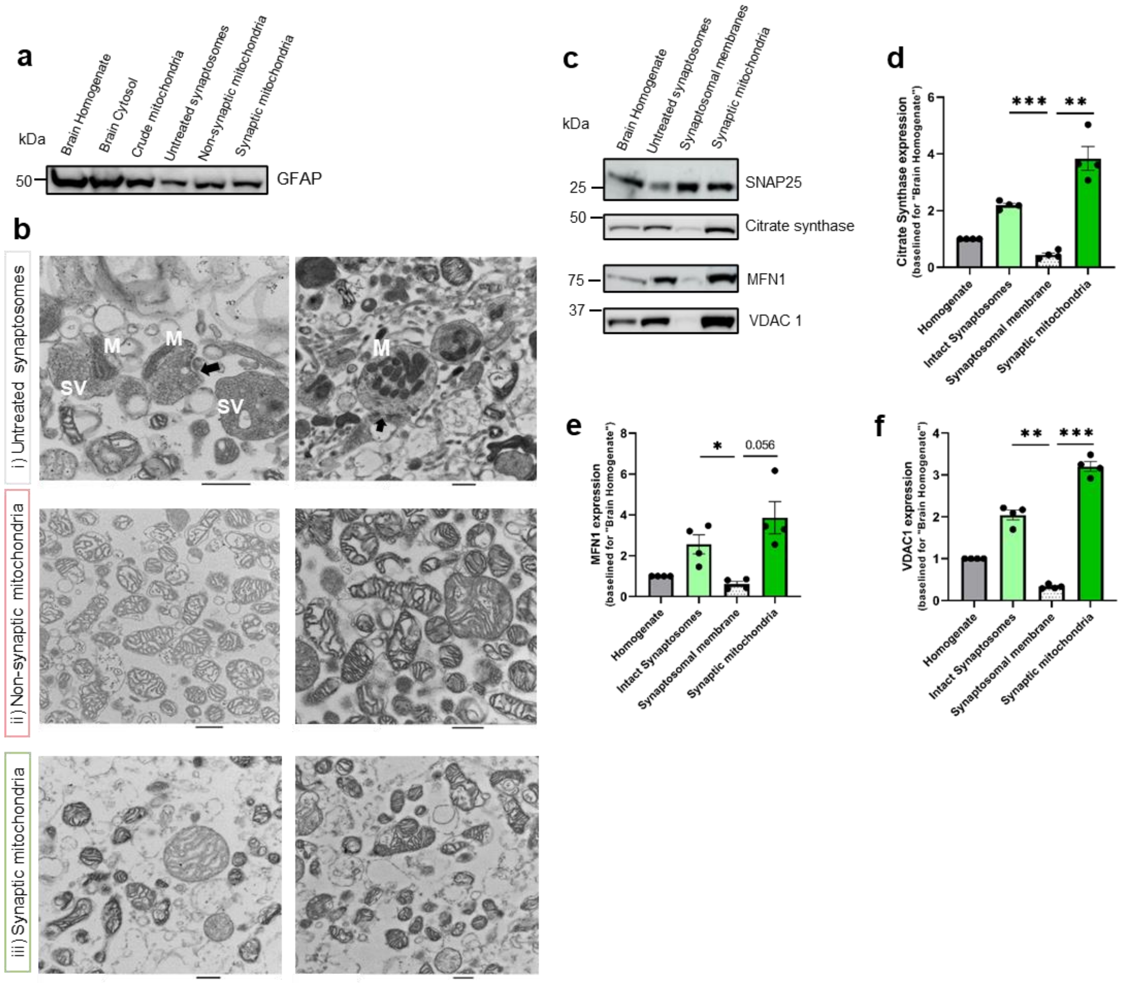


**Extended Data Fig. 1| a**, GFAP expression (**a**), an astrocytic marker in the fractions from brain isolation protocol (Fig. 1a) (n=2). TEM images of freshly isolated untreated synaptossomes; non-synaptic and synaptic mitochondria (related to Fig. 1); synaptic vesicles (SV), mitochondria (M) and post-synaptic density (black arrow). Scale bar = 500 nm. Expression (**c**) of SNAP25, a synaptosomal marker not membrane-associated and mitochondrial protein markers: Citrate Synthase (CS) (**c**, **d**), a mitochondrial matrix protein, Mitofusin1 (MFN1) (**c**, **e**) and VDAC1 (**c**, **f**), both outer mitochondrial membrane proteins in the fractions from brain isolation protocol (Fig. 1a) (n=4). **d**, **e** and **f** show means ± S.E.M., determined by one-way ANOVA with post-hoc Tukey test, *$P < 0.05$, **$P < 0.01$, ***$P < 0.001$.

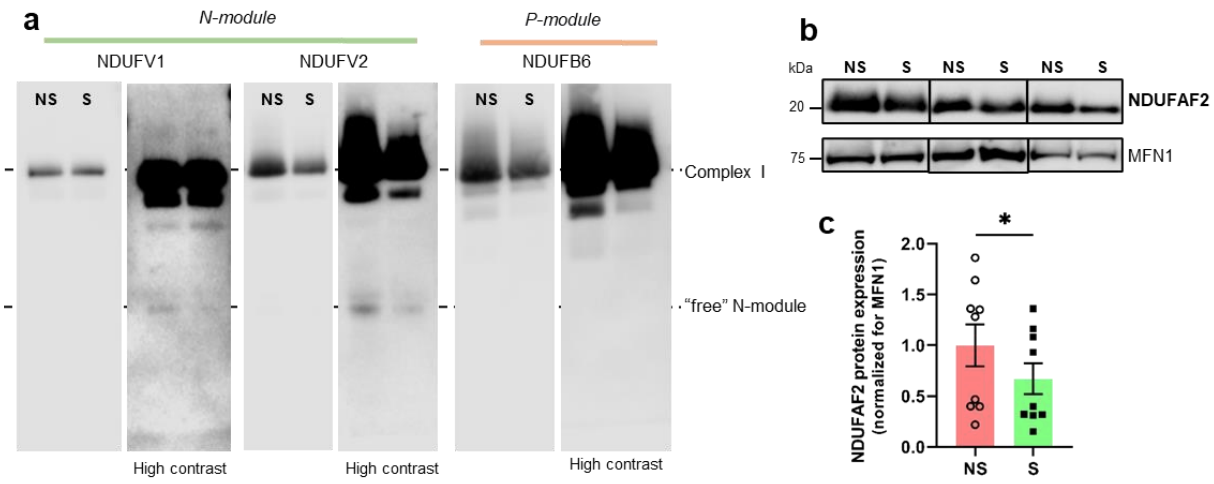


**Extended Data Fig. 2| N-module is not stalled in synaptic mitochondria.**

BN-PAGE (**a**) with DDM-treated mitochondrial fractions of synaptic (S) and non-synaptic (NS) mitochondria and probed for NDUFV1, NDUFV2 and NDUFB6 (n=2-3). Protein expression (**b**) and quantification (**c**) of NDUFAF2 expression in synaptic (S) and non-synaptic (NS) mitochondria. Bands intensity were normalized for Mitofusin1 (MFN1) expression. **c** show means ± S.E.M., determined by two-tailed paired t-test, $^{*}P < 0.05$ (n=9).
